# Transmission distortion and genetic incompatibilities between alleles in a multigenerational mouse advanced intercross line

**DOI:** 10.1101/2021.06.09.447720

**Authors:** Danny Arends, Stefan Kärst, Sebastian Heise, Paula Korkuc, Deike Hesse, Gudrun A. Brockmann

**Author notes:** To whom correspondence should be addressed. *Address for correspondence* Gudrun A. Brockmann.

## Abstract

**Background/Objectives:** While direct additive and dominance effects on complex traits have been mapped repeatedly, additional genetic factors contributing to the heterogeneity of complex traits have been scarcely investigated. To assess genetic background effects, we investigated transmission ratio distortions (TRDs) of alleles from parent to offspring using an advanced intercross line (AIL) of an initial cross between the mouse inbred strains C57BL/6NCrl (B6N) and BFMI860-12 (BFMI).

**Subjects/Methods:** 341 males of generation 28 and their respective 61 parents and 66 grandparents were genotyped using Mega Mouse Universal Genotyping Arrays (MegaMUGA). TRDs were investigated using allele transmission asymmetry tests, and pathway overrepresentation analysis was performed. Sequencing data was used to test for overrepresentation of non-synonymous SNPs in TRD regions. Genetic incompatibilities were tested using the Bateson-Dobzhansky-Muller two-locus model.

**Results:** 62 TRD regions were detected, many in close proximity to the telocentric centromere. TRD regions contained 44.5% more non-synonymous SNPs than randomly selected regions (182 vs. 125.9 ± 17.0, P < 1×10^−4^). Testing for genetic incompatibilities between TRD regions identified 29 genome-wide significant incompatibilities between TRD regions (P_(BF)_ < 0.05). Pathway overrepresentation analysis of genes in TRD regions showed that DNA methylation, epigenetic regulation of RNA, and meiotic/meiosis regulation pathways were affected independent of the parental origin of the TRD. Paternal BFMI TRD regions showed overrepresentation in the small interfering RNA (siRNA) biogenesis and in the metabolism of lipids and lipoproteins. Maternal B6N TRD regions harbored genes involved in meiotic recombination, cell death, and apoptosis pathways. The analysis of genes in TRD regions suggests the potential distortion of protein-protein interactions influencing obesity and diabetic retinopathy as a result of disadvantageous combinations of allelic variants in *Aass*, *Pgx6* and *Nme8*.

**Conclusions:** Using an AIL significantly improves the resolution at which we can investigate TRD. Our analysis implicates distortion of protein-protein interactions as well as meiotic drive as the underlying mechanisms leading to the observed TRD in our AIL. Furthermore, genes with large amounts of non-synonymous SNPs located in TRD regions are more likely to be involved in pathways that are related to the phenotypic differences between the parental strains. Genes in these TRD regions provide new targets for investigating genetic adaptation, protein-protein interactions, and determinants of complex traits such as obesity.

## Introduction

Over the last two decades genome wide association studies (GWAS) have identified a virtual avalanche of genetic variants associated with complex phenotypes and diseases (Buniello et al. 2019). However, although numerous, these identified genetic variations only partially explain the observed heritability in complex phenotypes either individually or combined (Zuk et al. 2012). This so called “missing heritability” problem has been observed for many complex phenotypes and diseases, such as obesity which has long been known to be a major risk factor for many diseases in the later course of life (Kopelman 2007; Tremmel et al. 2017). For obesity, as well as other complex phenotypes, a great amount of effort was spent finding genetic determinants (Speliotes et al. 2010; Willer et al. 2009). As a result, a large number of genetic variants contributing to obesity have been identified (https://www.genome.gov/gwastudies). However, most loci found by GWAS had small effects (Willer et al. 2009; Shungin et al. 2015). For example, the 97 significant loci identified for body mass index (BMI), accounted for only 2.7% of the corresponding variation (Locke et al. 2015).

One of several reasons discussed for the “missing heritability” problem in GWAS is the genetic heterogeneity of loci contributing to complex traits among the individuals in populations (Shungin et al. 2015; Heid et al. 2010). In addition to direct genetic effects, substantial phenotypic variation among individuals can be caused by preferential allele combinations or by allele incompatibilities in the genome of individuals. Allele incompatibilities are combinations of two (or more) alleles which when inherited together cause a disadvantage for the individual. Inheriting the disadvantageous allele combination leads to a survival disadvantage for this individual (e.g. less vigor, less successful reproduction). Allelic interactions leading to such disadvantages can be detected as transmission ratio distortion (TRD) from parent to offspring (also called allele transmission bias). Although TRD has been widely reported in a wide range of species (Huang et al. 2013b; Lyon 2003; Li et al. 2019), its functional impact on complex phenotypes has rarely been studied.

To investigate genetic background effects such as TRD, we examined three generations (26, 27, and 28) of an advanced intercross line (AIL) between two inbred mouse lines, the Berlin Fat Mouse Inbred (BFMI) line and C57BL/6NCrl (B6N) line. In AIL populations from two inbred founders, only two parental alleles can segregate at each locus, making the population heterogeneous but less complex than human populations. Therefore, an AIL population is well suited to study deviations from Mendelian inheritance.

The BFMI is an inbred line generated from an outbred population descending from several different founder mice bought at pet shops across Berlin. The BFMI line had been selected for high fatness for more than 100 generations before it was inbred (Wagener et al. 2006). Unfortunately, the original founders of the BFMI do not exist anymore. However, whole genome DNA sequencing data showed that the BFMI genome is a mixture of *Mus musculus musculus* and *Mus musculus domesticus*.

Recently, a major recessive mutation responsible for the juvenile obesity phenotype (*jObes1*) in BFMI mice was fine-mapped to a 370 kb region on chromosome 3 (Arends et al. 2016). Complementation tests suggested *Bbs7* as the most likely causal gene in this region. The *jObes1* locus accounted for around 40% of the body weight variance (Neuschl et al. 2010; Arends et al. 2016), while environmental effects accounted for 34%. Hence, around 26% of the variance in body weight is still unexplained.

During the process of long-term selection for a phenotype (e.g. high fatness), enrichment or even fixation of alleles that positively contributed to the selection response have been observed (Hirsch et al. 2014). Complimentary, the frequency of conflicting alleles impairing fitness, survival of gametes or embryos would be expected to be reduced or lost in the process of selection. In particular in complex traits, such as obesity, where many genes with diverse allelic variants contribute to the phenotype, the compatibility of interacting alleles is expected to be a driving force for the selection response. Therefore, long-term selection can be considered as co-evolution of alleles during the process of adaptation to selection pressure, environment, and genetic background.

The same principles of shaping the genomic composition occurred during the inbreeding history of every inbred mouse line, including BFMI and B6N, which were used in our experiment. Experimental inbreeding usually starts with several full-sib families (Flurkey et al. 2009). During the process of repeated mating of full-sibs, when the genome gets more and more reduced to one haplotype, some inbred families go extinct because of direct lethal recessive allele effects, lethal combinations of alleles across the genome, or the inbred family collapses because of low vigor or insufficient reproduction (Whitlock 2000; Fitzpatrick and Evans 2009; Zajitschek et al. 2009) eventually as a result of genomic incompatibility. However, low level incompatibilities, which do not directly cause lethality or affect fertility might be retained, invisible, inside an inbred line. During the inbreeding process haplotypes get reduced, and incompatibilities might survive since there is no choice of alternative allele anymore. In the end, a kind of optimized genome remains alive as established inbred strain.

## Results

### Allele transmission ratios from heterozygous parents to offspring

The probability for the transmission of parental alleles to their offspring can be calculated according to Mendelian laws. Deviations from those expected inheritance patterns might have genetic reasons that we intend to identify. To test for TRDs, we used all 341 males of generation 28 of the AIL and tested how their parents (generation 27) transmitted their alleles to this generation. TRD was detected for 62 genomic regions at a genome-wide Bonferroni corrected significance level of 0.01. These regions can be grouped by the preferentially transmitted allele based on the parental origin (paternal / maternal) and the founder strain origin (B6N / BFMI). Significant paternal allele TRD was detected for 1,068 out of 18,114 tested SNPs. Paternally affected TRD of SNPs clustered into 31 chromosomal regions, due to linkage between neighboring SNPs. For maternal TRD, 1,138 SNPs located in 31 regions were found (Supplemental_Table_S1). Overlaying the paternal and maternal TRD regions showed that 14 regions showed both paternal and maternal TRD. In overlapping TRD regions always the same founder allele of either mouse strain B6N or BFMI was preferentially transmitted.

TRD was detected consistently across large genomic regions, in which a high number of markers showed the same transmission bias for one of the two founder alleles B6N or BFMI. The 19 regions showing TRD supported by at least 50 markers are shown in Table 1, all detected TRD regions and their observed transmission distortions are summarized in Supplemental_Table_S1, and visualized in Figure

**Table 1:** Transmission ratio distortion from paternal and maternal side, supported by at least 50 SNPs (p < 0.01) per identified region. Region ID = the identifier of the region Pat_R# stands for paternal transmission ratio distortion and Mat_R# stands for maternal transmission ratio distortion within the region, Chr = Chromosome on which the distortion was detected. Proximal, Top, and Distal = start, top, and end positions of the region on the chromosome (based on the GRCm38.p6 / mm10 genome), preferred allele = the allele preferentially transmitted, nSNPs = number of SNP markers on the array that support the TRD region, average transmission counts for founder alleles across all markers in the region are listed with their standard deviation in brackets. Top marker as well as transmission at the top marker and BFMI allele % distortion is shown in the last 4 columns. See Supplemental_Table_S1 for an overview of all 31 paternal and 31 maternal regions. The observed very small standard deviations in almost all regions indicate that the distortion observed is consistent across the regions identified.

| RegionID | Chr | Proximal | Top | Distal | RegionSize | nSNPs | Preferred Allele | Transmissions regionaverage (SD) |  | Topmarker | Transmission Topmarker |  | BFMIallele %distortion |
| --- | --- | --- | --- | --- | --- | --- | --- | --- | --- | --- | --- | --- | --- |
|  |  |  |  |  |  |  |  | BFMI | B6N |  | BFMI | B6N |  |
| Pat_R1 | 1 | 3,668,628 | 11,643,615 | 14,698,538 | 11,029,910 | 68 | BFMI | 146.1 (16.3) | 41.1 (6.6) | UNC109624 | 152 | 30 | 67.0 |
| Pat_R3 | 4 | 3,569,913 | 6,093,982 | 12,555,306 | 8,985,393 | 83 | BFMI | 168.0 (5.0) | 55.3 (3.9) | UNC6664886 | 169 | 48 | 55.8 |
| Pat_R7 | 4 | 76,193,199 | 78,074,351 | 81,245,565 | 5,052,366 | 58 | BFMI | 79.6 (9.0) | 15.8 (3.0) | UNC7556251 | 84 | 15 | 69.7 |
| Pat_R17 | 12 | 5,253,913 | 8,416,509 | 12,177,986 | 6,924,073 | 92 | BFMI | 163.3 (28) | 53.5 (9.6) | UNC20594325 | 207 | 47 | 63.0 |
| Pat_R21 | 16 | 5,617,528 | 14,151,479 | 14,151,479 | 8,533,951 | 93 | B6N | 31.5 (7.2) | 147.9 (15.1) | UNC26373573 | 35 | 177 | -67.0 |
| Pat_R22 | 17 | 3,264,958 | 5,991,544 | 11,318,508 | 8,053,550 | 77 | B6N | 34.0 (16.2) | 99.4 (27.1) | UNC170286629 | 16 | 139 | -79.4 |
| Pat_R23 | 17 | 47,490,686 | 48,644,966 | 50,134,302 | 2,643,616 | 50 | B6N | 41.7 (30.8) | 117.1 (10.1) | UNC27963988 | 19 | 110 | -70.5 |
| Pat_R24 | 18 | 4,516,519 | 12,242,865 | 13,902,153 | 9,385,634 | 71 | BFMI | 158.7 (2.9) | 61.4 (3.3) | UNC28742422 | 155 | 56 | 46.9 |
| Pat_R25 | 18 | 38,281,545 | 38,975,190 | 42,987,483 | 4,705,938 | 54 | BFMI | 168.7 (4.1) | 54.2 (9.3) | UNC29080241 | 168 | 46 | 57.0 |
| Mat_R1 | 1 | 3,668,628 | 3,668,628 | 21,099,704 | 17,431,076 | 94 | BFMI | 136.2 (17.8) | 56.9 (10.3) | UNC010515443 | 178 | 69 | 44.1 |
| Mat_R2 | 3 | 6,274,425 | 11,119,314 | 17,230,305 | 10,955,880 | 53 | B6N | 24.9 (13.5) | 129.2 (5.8) | backupUNC030002827 | 13 | 126 | -81.3 |
| Mat_R5 | 3 | 133,540,430 | 136,529,917 | 137,921,098 | 4,380,668 | 52 | B6N | 35.4 (4.1) | 105.3 (12.8) | UNC6326383 | 33 | 113 | -54.8 |
| Mat_R6 | 4 | 3,569,913 | 3,918,966 | 12,555,306 | 8,985,393 | 83 | BFMI | 144.5 (5.5) | 50.1 (6.4) | UNC6640040 | 135 | 38 | 56.1 |
| Mat_R12 | 6 | 48,276,599 | 48,276,599 | 53,750,577 | 5,473,978 | 53 | B6N | 24.8 (5.0) | 80.5 (15.4) | UNC11030573 | 27 | 96 | -56.1 |
| Mat_R15 | 9 | 67,022,716 | 74,576,994 | 75,773,843 | 8,751,127 | 54 | B6N | 37.3 (12.9) | 110.8 (16.1) | UNC090145124 | 23 | 99 | -62.3 |
| Mat_R16 | 9 | 106,017,492 | 106,605,721 | 107,516,430 | 1,498,938 | 58 | B6N | 29.3 (4.5) | 87.1 (2.4) | UNC17077906 | 27 | 87 | -52.6 |
| Mat_R23 | 16 | 5,617,528 | 5,617,528 | 18,450,764 | 12,833,236 | 163 | B6N | 23.5 (8.7) | 108.4 (11.2) | UNC160000883 | 10 | 130 | -85.7 |
| Mat_R25 | 17 | 3,264,958 | 5,991,544 | 11,318,508 | 8,053,550 | 77 | B6N | 20.0 (4.4) | 106.8 (22.4) | UNC170286629 | 20 | 164 | -78.3 |
| Mat_R28 | 18 | 4,516,519 | 8,317,246 | 15,000,978 | 10,484,459 | 81 | BFMI | 170.6 (19.5) | 52.0 (2.8) | UNC28690832 | 50 | 197 | -59.5 |

1. Genotypes, genetic map, and pedigree of the AIL individuals can be found in Supplemental_Table_S2.

As an example, Pat_R3 is a region on chromosome 4 from 3.5 to 12.5 Mb, which showed paternal TRD of the BFMI allele supported by 83 markers. In the AIL population, we observed at each marker around 217 paternal allele transmissions from generation 27 to 28, meaning we expect 108.5 transmissions of the BFMI, as well as 108.5 transmissions of the B6N allele. However, in this region markers on average showed 168 (+/− 5.0) transmissions of the BFMI allele from heterozygous fathers to their offspring, while the B6N allele was only transmitted 55.3 (+/− 3.9) times. At the top marker in this region on chromosome 4 (UNC6664886), we observed 169 BFMI versus 48 B6N transmissions. This means that transmission of the BFMI allele was observed 55.8 % more often than expected by Mendelian inheritance, the likelihood of this happening was estimated by χ^2^-test to be lower than 1×10^−14^.

When performing the same tests for the allele transmission from AIL generation 26 to 27 (transmission from grandparents (n = 66) to parents (n = 61)), where sample sizes were much smaller, we relaxed our threshold for significance to p < 0.05. In generation 26 to 27, 0 and 38 SNPs still showed significant paternal or maternal TRD, respectively. The overlap between SNPs detected in generations 26 to 27 versus 27 to 28 was 100%, meaning that all TRD seen from generation 26 to 27 was also found (much more significant) from generations 27 to 28.

In our population, we observed that the TRD is a local event affecting many SNPs in a well-defined chromosomal region due to linkage between neighboring SNPs. Since, many recombinations have accumulated over 28 generations of mating, the length of the TRD affected regions is between 41.1 kb and 17.4 Mb (Supplemental_Table_1). In these regions, SNPs that showed TRD are tightly linked (Figure 1), which can be seen by TRD SNPs clustering together into regions. Very small standard deviations of averaged TRD transmissions (Table 1) were observed for all regions, which indicates that distortion observed was consistent across the TRD region. This was further supported by the observation that all SNPs in a certain region always showed the same direction of transmission towards one of the alleles from the original founder strains of the AIL population. For example, Mat_R1 showed the BFMI allele was preferentially transmitted in the maternally distorted region on chromosome 1 between 3.7 and 21.1 Mb. For the 94 markers in this region, we observed 136.2 +/− 17.8 transmissions of the BFMI allele versus 56.9 +/− 10.3 transmissions of the B6N allele from mother generation 27 to offspring (generation 28).

**Figure 1:**
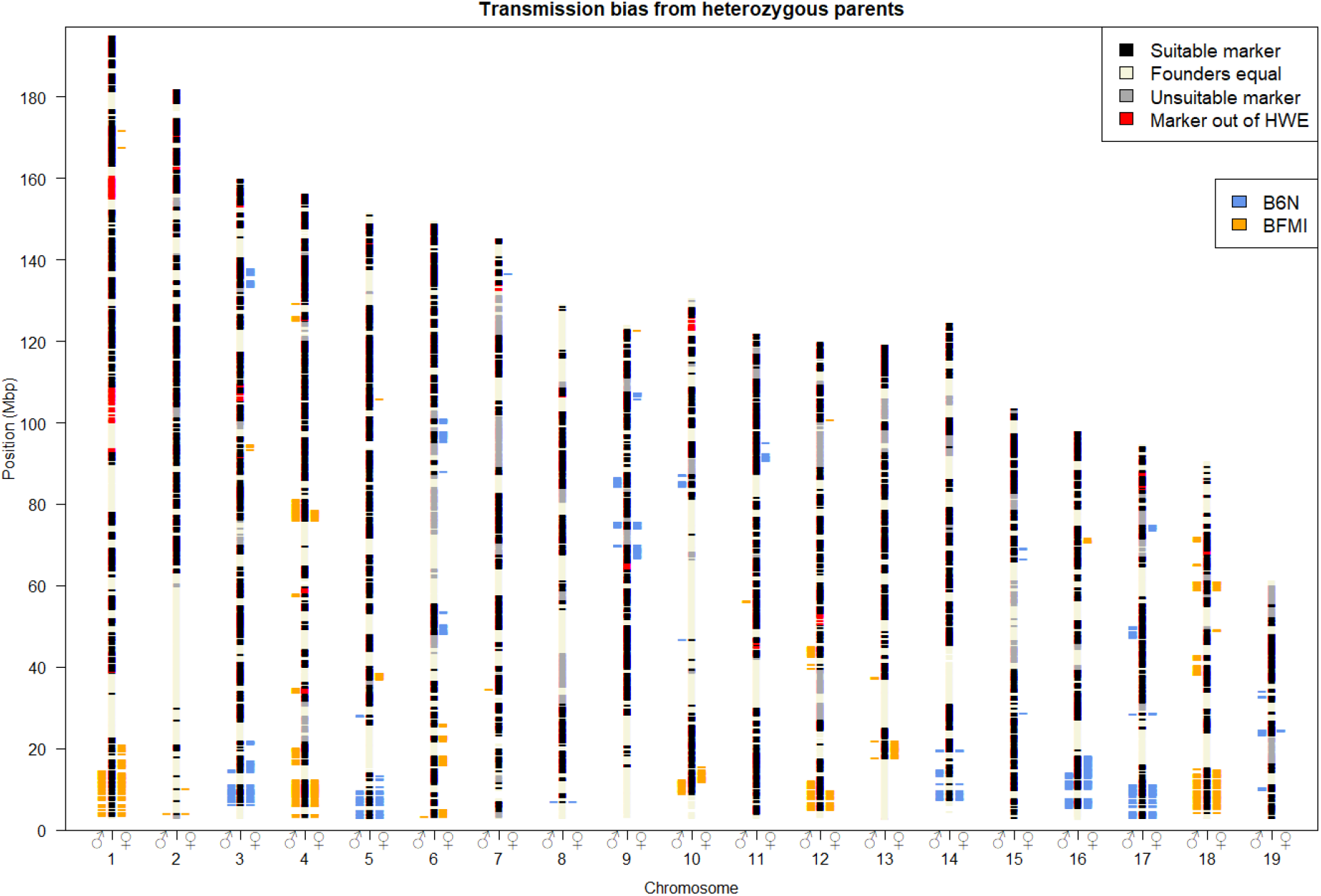
Genomic regions showing allele transmission ratio distortion towards generation 28. Bars left of the chromosomes mark the SNPs which show paternal TRD (♂); bars on the right side show maternal TRD (♀), using a genome wide significance level of P < 0.01. Colors show the origin of the allele preferentially transmitted, blue: B6N allele, orange: BFMI allele. Chromosomal black areas (suitable markers) contain markers which passed quality control steps, segregate between the founder lines (BFMI and B6N), and have at least ten heterozygous parents in generation 27 required to perform a valid X2 test. Chromosomal beige areas (Founders equal) are markers at which the BFMI and B6N have the same allele, these markers do not segregate in the AIL population, and cannot be tested for TRD. Chromosomal gray areas (unsuitable markers) have not been tested due to lack of heterozygous parents in generation 27 at these markers. Chromosomal red areas are not in Hardy-Weinberg equilibrium (HWE) in generation 28, since HWE is an assumption underlying a valid TRD test, these areas were excluded from TRD analysis.

Interestingly, regions showing significant TRD on 10 out of 19 autosomes (autosomes 1, 3, 4, 5, 10, 12, 14, 16, 17, 18) are located close to the telocentric centromere (Figure 1). These 10 telocentric centromere regions showed both paternal and maternal TRD with a consistent preference of the founder allele. While these telocentric centromere regions showed TRD for both paternal and maternal alleles, we observed that non-telocentric centromere regions (e.g. Mat_R12, Table 1) tend to show TRD only when inherited from either the paternal or the maternal side.

### Genetic variants in TRD regions

To identify candidate genes for each region, and to investigate possible causes for the observed TRD, protein coding genes located in TRD regions were examined. Sequence variants were detected by comparing the BFMI sequence to the B6N reference genome (ENSEMBL, GRCm38.p6) (Supplemental_Table_S3). In the 62 identified TRD regions, 1,167 unique protein coding genes were located. In detail, these were 292 genes in Pat_BFMI regions, 362 in Mat_BFMI, 335 in Pat_B6N, and 567 in Mat_B6N. Among those, 389 genes were overlapping between paternal and maternal TRD regions.

In the 1,167 unique protein coding genes located in 62 TRD regions, 182 non-synonymous SNPs (nsSNPs) were found located in 128 (10.9%) genes. Permutation analysis showed that the density of nsSNPs in the TRD regions was 1.445 times higher than expected from random distribution. Results from 50,000 permutations showed an average of 125.9 ± 17.0 (SD) nsSNPs per 1,167 randomly selected genes with a maximum value of 180 SNPs observed during permutation. These results provide evidence that nsSNPs are significantly overrepresented in TRD regions (P < 1×10^−4^).

### Pathway overrepresentation analysis

Pathways analysis was performed twice, once we investigated pathway overrepresentation by including all protein-coding genes in the specified TRD regions, followed by only investigating genes that carry nsSNPs.

Analysis of all genes in TRD regions with higher transmission of the paternal BFMI allele (Pat_BFMI) showed slight but significant pathway overrepresentation of the three pathways “Post-transcriptional silencing by small RNAs” (P_(BH)_ = 0.008), “Small interfering RNA (siRNA) biogenesis” (P_(BH)_ = 0.011), and “MicroRNA (miRNA) biogenesis” (P_(BH)_ = 0.016) (Supplemental_Table_S4 - Pat_BFMI for the full list).

All genes located in maternal inherited (Mat_BFMI) showed highly significant overrepresentation for pathways such as “DNA methylation” (P_(BH)_ < 5.7×10^−16^), “Meiotic Recombination” (P_(BH)_ < 6.61×10^−15^), “Packaging Of Telomere Ends” (P_(BH)_ < 1.70×10^−13^), “Chromatin organization” (P_(BH)_ < 2.66×10^−10^) and “Deposition of new CENPA-containing nucleosomes at the centromere” (P_(BH)_ < 3.67×10^−10^) (Supplemental_Table_S4 - Mat_BFMI for the full list). All these pathways are involved in chromosome stability/maintenance as well as centromere and nucleosome organization. We also found strong overrepresentation of the “Signaling by Wnt” (P_(BH)_ = 1.09×10^−5^) pathway, as well a weak overrepresentation of the “Retinol metabolism” (P_(BH)_ = 0.010) pathway. These are two interesting pathways in the context of BFMI mice, which will be elaborated in more detail in the discussion section.

No strong overrepresentation or overlap was found for all genes located in paternal/maternal B6N TRD regions (Supplemental_Table_S4 - Pat_B6N & Mat_B6N). Only three pathways were found weakly overrepresented when using all genes from maternal B6N (Mat_B6N) TRD regions: “Nitrogen metabolism” (P_(BH)_ < 0.019), “Reversible hydration of carbon dioxide” (P_(BH)_ < 0.033), and “Osteoclast differentiation” (P_(BH)_ < 0.042).

If we focused on genes with nsSNPs in TRD regions, where the BFMI allele is preferentially passed by the father (Supplemental_Table_S4 - Pat_BFMI_SNP), no strong pathway overrepresentation was observed. Only two pathways reach significance after Benjamini-Hochberg correction: “S Phase” (P_(BH)_ = 0.015) and “Extracellular matrix organization” (P_(BH)_ = 0.025). However, the number of genes found in TRD regions (2 for both) versus the total number of genes annotated to these pathways make this overrepresentation very weak (117 and 216 respectively). If we examined the genes with nsSNPs from TRD regions of paternal B6N allele transmission (Supplemental_Table_S4 - Pat_B6N_SNP), “Interferon Signaling” (P_(BH)_ = 0.016), “Cell cycle” (P_(BH)_ = 0.023), and “Metabolism of lipids and lipoproteins” (P_(BH)_ = 0.023) were weakly significantly overrepresented. Again, the numbers of genes in TRD regions is small compared to the total number of genes annotated to these pathways.

Genes with nsSNPs from TRD regions of the maternal BFMI allele (Supplemental_Table_S4 - MAT_BFMI_SNP) showed only one very weak significant pathway overrepresentation: “Extracellular matrix organization” (P_(BH)_ = 0.046), which was also found for genes with nsSNPs in TRD regions where the BFMI allele was preferentially passed by the father (Pat_BFMI_SNP). Genes with nsSNPs in TRD regions of the maternal B6N allele (Supplemental_Table_S4 - Mat_B6N_SNP) also showed overrepresentation of multiple pathways overlapping with pathways found for genes with nsSNPs in paternal B6N regions (“Cell cycle”, “Interferon Signaling”, “Metabolism”, and “Metabolism of lipids and lipoproteins”). Furthermore, maternal genes with nsSNPs in TRD regions contributed also to cell death and apoptosis (pathways “Cell death signalling via NRAGE, NRIF and NADE” (P_(BH)_ = 0.04) and “p75 NTR receptor-mediated signaling” (P_(BH)_ = 0.049)).

### Genetic incompatibilities

Since functional inaptitude of alleles of interacting pairs of genes could be causal for TRD, we searched for evidence of genetic incompatibilities by a pairwise search between all 62 TRD regions against each other using the Bateson-Dobzhansky-Muller model. This test identifies outliers (significant deviation from Mendelian expectation) between allele pairs by investigating allele frequencies. Severe deviations from Mendelian expectation is interpreted as resulting from negative epistatic interactions between incompatible loci. This search identified genome-wide significant pairwise incompatibilities (P_(BF)_ < 0.05) for 29 out of 62 TRD regions (Figure 2), of which 19 were classified as highly significant (P_(BF)_ < 0.01). The high number of TRD regions (46.8 %) showing evidence for one or more genetic incompatibilities suggests that genetic incompatibilities are an important contributor to TRD.

**Figure 2:**
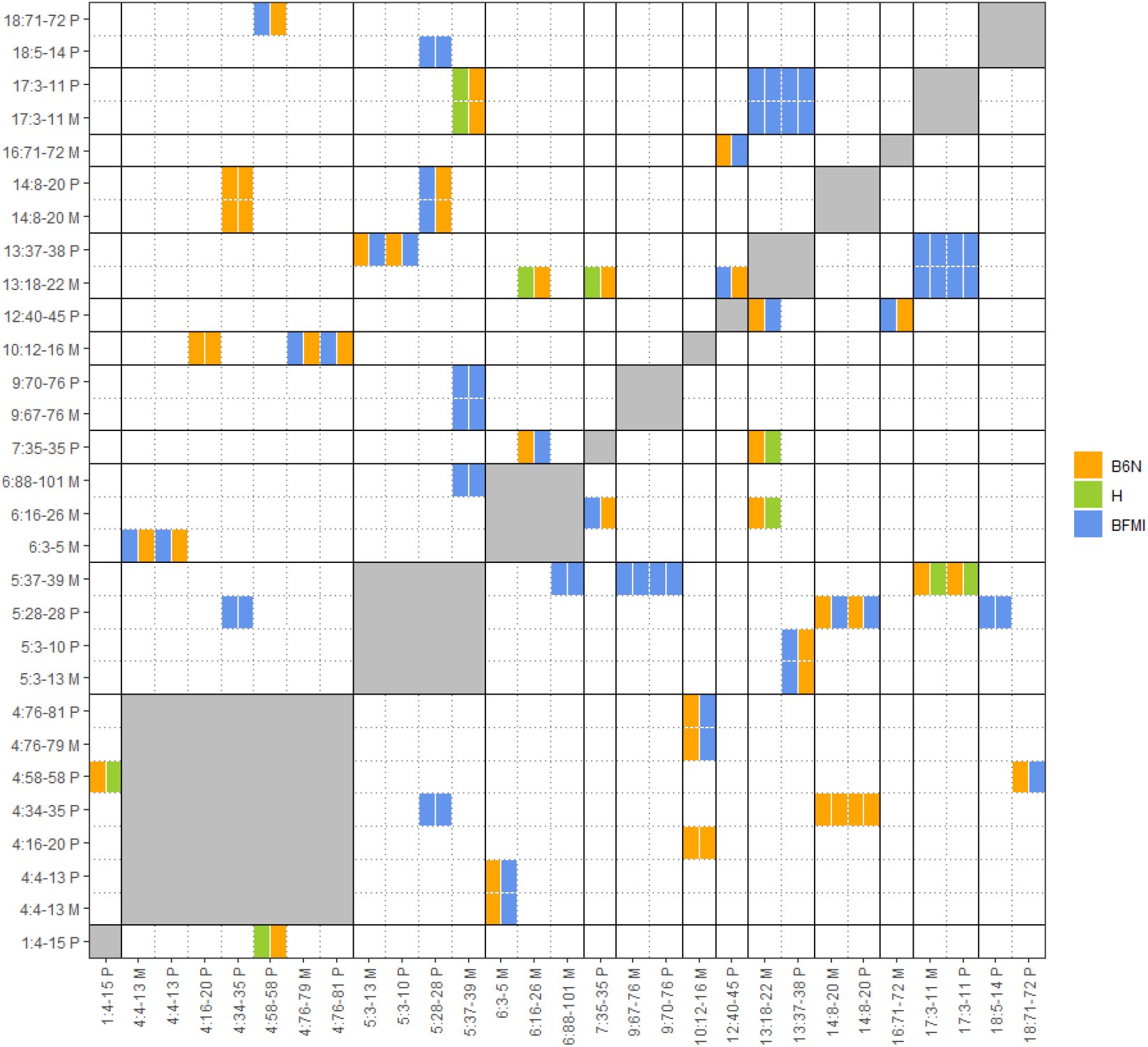
Significant genetic incompatibilities between regions showing TRD. Heat map showing the pairwise genetic incompatibility scan between TRD regions, genome-wide P_(BF)_ < 0.05. The allele combination (M1|M2) which is most reduced (in percentages) between the observed and expected allele combinations are shown in the figure with colors denoting the founder allele combination M1 (x-axis) and M2 (y-axis). Names of regions are composed of chr:start-end allele origin; start and end positions are given in megabase pairs; furthermore, the TRD origin is coded by M for maternal and P for paternal. When two regions were located on the same chromosome the genetic incompatibility test was not performed (gray areas), since the pairwise genetic incompatibility test can only be performed on loci which are not in linkage.

Analysis of protein-protein interactions between all 128 protein-coding genes with nsSNPs located in TRD regions showed 331 known interactions between the protein products of these genes. When we ignored genes located on the same chromosome (for which genetic incompatibility tests cannot be performed) we ended up with 273 known protein-protein interactions.

Within the 29 regions that showed genome-wide evidence (P_(BF)_ < 0.05) for genetic incompatibilities, 44 out of the 128 (34.4%) protein-coding genes with nsSNPs reside. In total, five known physical protein-protein interactions exist in the Search Tool for the Retrieval of Interacting Genes/Proteins (STRING) database between these 44 genes.

Within these known protein-protein interactions, we found an interaction between alpha-aminoadipic semialdehyde synthase (Aass) located in Mat_R11 (6:16-26 M) with glutathione peroxidase 6 (Pgx6) as well as NME/NM23 Family Member 8 (Nme8) which are both located in Mat_R20 (13:18-22 M). All three of these genes are interesting, given that all three genes are involved in obesity and/or diabetic retinopathy, which are the obvious phenotypic difference between the founders inbred strains BFMI and B6N. The Aass protein is involved in the major mitochondrial lysine degradation pathway (Papes et al. 1999; Sacksteder et al. 2000) and was found to be downregulated in obese compared with lean co-twins (Heinonen et al. 2015). With regard to Pgx6, glutathione peroxidase activity is suppressed in diabetic compared to healthy controls, with a more pronounce suppression in obese compared to nonobese diabetics (Singhai et al. 2011). Glutathione peroxidase activity was found associated with diabetic retinopathy (Rodríguez-Carrizalez et al. 2014). Nme8, encodes an axoneme protein, and mutations in the *Nme8* gene have been implicated to cause primary ciliary dyskinesia (Duriez et al. 2007). Furthermore, the genetic region in which the *Nme8* gene is located was identified in human GWAS as a locus which might be involved in childhood obesity in the Hispanic population (Comuzzie et al. 2012). *Nme8* is a very interesting gene to come up during this analysis, because of its relation to primary ciliary dyskinesia. The *Bbs7* gene was previously identified as the most likely causal gene for the obesity phenotype of the BFMI mouse (Arends et al. 2016). The Bbs7 protein is part of the BBSome complex which is a heterooctameric protein complex that plays a central role in primary cilia homeostasis (Klink et al. 2020).

Furthermore, a protein-protein interaction between acyloxyacyl hydrolase (Aoah), located Mat_R20 (13:18-22 M) and protein tyrosine phosphatase receptor type Z1 (*Ptprz1*) Mat_R11 (6:16-26 M) was found. Aoah is a lipase that plays an important role in the defense against gram-negative bacterial infection (Lu et al. 2008). Aoah^−/−^ mice on a high-carbohydrate diet develop non-alcoholic steatohepatitis (NASH) and both serum triglyceride and cholesterol were found significantly increased (Ojogun 2008). The *Ptprz1* gene is annotated to the insulin receptor recycling pathway, and protein tyrosine phosphatases (PTPs) are required for the dephosphorylation of the insulin receptor (Fischer et al. 1991). Again, both genes identified by our approach are clear candidate genes when considering the major phenotypic differences of the founder strains BFMI and B6N.

The next identified protein-protein interaction between genes with nsSNPs in TRD regions showing genetic incompatibilities was between myosin IE (Myo1e), located Mat_R15 (9:67-76 M) and serine/threonine kinase 32B (Stk32b) Mat_R9 (5:37-39 M). Myo1e is part of the nonmuscle class I myosins which are a subgroup of the unconventional myosin protein family and function as actin-based molecular motor. The *Stk32b* gene is annotated into the “Sweet Taste Signaling” pathway (GeneCards Human Gene Database 2021), and deletion of the gene was associated with Ellis-Van Creveld Syndrome (Temtamy et al. 2008) in humans. This interaction doesn’t have a clear link to the phenotypic differences between BFMI and B6N.

Moreover, protein-protein interaction was detected between myosin VC (Myo5c), located Mat_R15 (9:67-76 M) and solute carrier family 2 member 9 (Slc2a9) located in Mat_R9 (5:37-39 M). The Myo5c protein is involved in actin-based membrane trafficking in many physiologically crucial tissues. In humans (and mice), Myo5c is particularly abundant in epithelial and glandular tissues such as: pancreas, prostate, mammary, stomach, colon, and lung (Rodriguez and Cheney 2002). *Myo5c* knockout mice show a decrease in total body fat amount and an increased lean body weight (Blake et al. 2021; Mouse Genome Database (MGD) 2021). It’s protein interaction partner Glut9 (*Slc2a9*) is part of the SLC2A facilitative glucose transporter family. Members of this family play a role in maintaining glucose homeostasis. Glut9 does not transport glucose, but is classified as a urate transporter. Mutations in the *Slc2a9* gene have been shown to be causal for renal hypouricemia (Matsuo et al. 2008; Dinour et al. 2010), mice lacking the Glut9 protein show early onset metabolic syndrome (DeBosch et al. 2014).

These known protein-protein interactions between genes in TRD regions with nsSNPs, lead us to hypothesize that disturbed protein-protein interactions resulting from amino acids changes due to nsSNPs within several proteins of a protein complex are likely one of the driving forces causing the TRD observed in the BFMIxB6N advanced intercross line.

## Conclusions and discussion

In this study, we examined an AIL population originating from a cross between the obese mouse line BFMI and the standard mouse line B6N in generations 26 to 28 for TRD from parents and grandparents to offspring. The most significant finding of this study was the detection of 62 genomic regions showing TRD in the genotype data from generations 27 to 28.

We considered three possible explanations for the widespread TRD we observe in our AIL, (1) independent selection at each locus, (2) gametic or meiotic drive, and (3) preferential selection of combinations of alleles at two or more loci.

The first hypothesis that selection happens at each locus independently will often lead to nonproductive crosses and/or massive lethality after birth (Huang et al. 2013a). The argument against this first hypothesis is that the litter size in generation 28 does not deviate from the litter size in the parental inbred lines BFMI and B6N (data not shown). When incompatibilities are (embryonically) lethal this would cause a side-effect of significant TRD which should also be detectable as deviation from the Hardy Weinberg equilibrium (HWE) in the offspring generation (Paterson et al. 2009). Since regions out of HWE were excluded in our study, (embryonically) lethal alleles were not investigated in our AIL. This means that the observed TRD cannot be due to lethality, leading us to reject this hypothesis of direct independent selection at each locus as the cause for the TRD observed in our AIL.

The second possible mechanism for the TRD observed in this paper are the well-investigated examples in mouse from meiotic drive, such as the *t*-complex (Safronova and Chubykin 2013). In short, meiotic drive can be thought of as a conflict in which a selfish allele is able to use asymmetric meiosis in order to have a greater chance of being transmitted to the gamete. This mechanism in first instance fits our observations well, since peri-centromeric regions seem to be involved. Detected TRD regions in our AIL were observed located in close proximity to the telocentric centromere for 10 out of 19 autosomes. This observation is consistent with previous findings in e.g. *Drosophila,* where autosomal meiotic drivers, occur in heterochromatic regions around centromere and telomere (Brand et al. 2015). In mice, genetic incompatibilities in and around the centromeric regions between *Mus musculus musculus* and *Mus musculus domesticus* have been known for decades (Fel-Clair et al. 1998; Lenormand et al. 1997) and have been studied extensively in mouse populations near the hybrid zone (Teeter et al. 2008; Larson et al. 2018). Centromere strength differs between mouse strains and was found to predict the direction of meiotic drive in mice (Chmátal et al. 2014). Earlier findings showed no incompatibility between the chromosome 11 centromere region in hybrids between *Mus musculus musculus* and *Mus musculus domesticus* (Lanneluc et al. 2004). Our study confirms this finding, since we also did not observe TRD at the chromosome 11 centromeric region. Genome-wide DNA sequencing showed that BFMI is a hybrid between *Mus musculus musculus* and *Mus musculus domesticus* (data not shown, sequencing data available at SRA). As such, the AIL between BFMI and B6N might have revived incompatibilities stemming from meiotic drive between *musculus* and *domesticus* alleles. However, an argument against meiotic drive causing our TRD is that true meiotic drive would have led to fixation of the favored allele / haplotype within 26 to 28 generations (Kursel and Malik 2018). Since we do not observe this fixation, the meiotic drive hypothesis is unlikely to underly the widespread TRD observed in our AIL. Additionally, in mammals only female meiosis was found to be asymmetric (Brunet and Verlhac 2011; Kursel and Malik 2018), meaning that our observed paternal TRD is most likely not due to meiotic drive. However, we cannot exclude that this hypothesis might play a role for the maternal TRD regions observed near the centromeric regions. Furthermore, pathway overrepresentation analysis does show overrepresentation of pathways which point to meiotic drive in maternal BFMI TRD regions.

The third hypothesis is that TRD at each locus is not independent but rather caused by selection on preferential combinations of alleles or selection against detrimental allele combinations (Martin-DeLeon et al. 2005; Xie et al. 2019). nsSNPs in protein-coding genes located in TRD regions were investigated to see if this hypothesis could explain the TRD observed. In total, we found 182 nsSNPs in 128 genes within the 62 identified regions showing TRD. Based on permutation we would have expected to see only 125.9 ± 17.0 nsSNPs. The density of nsSNPs in genes in TRD regions was 44.5% higher than expected by chance. While the changes in amino acid sequence derived from nsSNPs in a single gene might not be sufficient to cause lethality or to reduce fitness, co-occurrence with SNPs in protein-protein interaction partners could cause such adverse effects, e.g. by affecting protein-protein binding leading to signaling problems (Xie et al. 2019). Such adverse effects could result in detectable TRD. On evolutionary time scales this is known as protein co-evolution, known to leave detectable footprints (Clark et al. 2011; Teppa et al. 2017). We hypothesize that this is what drives most TRD in our BFMI x B6N AIL. In our AIL we combine the genome of BFMI (a mix between *Mus musculus musculus* and *Mus musculus domesticus*) with B6N, a *Mus musculus musculus*. Our AIL as such forces together the evolutionary separated genomes of *Mus musculus musculus* and *Mus musculus domesticus* which might lead to resurging incompatibilities between proteins coded in different TRD regions.

However, it should be noted that in this paper we did not investigate the subspecific origin of the TRD regions and as such we can only hypothesize that protein-protein incompatibilities might be the underlying mechanism for the TRD we observe. Further research on the subspecific origin of the TRD would allow us to disentangle the contribution of each of the three hypotheses on our observed TRD.

Pairwise testing of genetic incompatibilities between the 62 identified TRD regions showed 29 genome-wide highly significant genetic incompatibilities in our AIL. Although our analysis shows that observed allele TRD is likely due to incompatibility between proteins in two or more TRD regions, genetic incompatibilities only account around half of TRD observed. Potentially, some incompatibilities could not be detected since (1) we limited our analyses to pairwise testing TRD regions, (2) incompatibilities might not always lead to detectable transmission ratio distortion, and (3) not all protein-protein interactions are known yet and/or stored in the STRING database. Furthermore, some genomic regions did not contain informative markers, and as such, they did not allow us to test for TRD in these regions. However, we cannot exclude meiotic drive for maternal TRD regions near the centromeres which might act alongside the genetic incompatibility hypothesis.

Pathways overrepresented in maternal BFMI TRD regions strongly point to meiotic drive with pathways such as: “DNA methylation”, “meiotic/meiosis regulation”, “Chromatin organization”, “Nucleosome assembly”, and “Telomere Maintenance” overrepresented. This is in line with the meiotic drive hypothesis being causal for some of the maternal TRD observed near the centromeres.

Paternal TRD regions showed overrepresentation of “Signaling by Wnt”, “Metabolism of lipids and lipoproteins”, and “Retinol metabolism”. These pathways point to incompatibilities, and genes located in TRD regions, such as acyl-CoA oxidase 2 (*Acox2*), fat binding proteins 4 and 5 (*Fabp4*, *Fabp5*), fatty acid desaturase 2 (*Fads2*) and malic enzyme 1 (*Me1*) which are known to be involved in energy partitioning and metabolism phenotypes in which the BFMI and B6N founders differ. Recent work on the retina of BFMI mice has shown differences in the rhodopsin layer of BFMI vs. B6N mice, pointing towards an impaired retina function in BFMI mice. Eyes of the BFMI showed definite characteristics of retinal degeneration in terms of a dysfunction of the rhodopsin transport and a reduction in the outer nuclear layer (ONL) thickness (Brockmann C. et al. 2017). This might explain why genes located in the “Retinol metabolism” pathway come up as significantly overrepresented. Our TRD analysis identifies genes within TRD regions that could be considered as possible candidate genes for retinal degeneration in mice and humans.

When looking into pathways that were overrepresented while analyzing TRD genes with nsSNPs, pathways such as “Cell Cycle”, “Metabolism of lipids and lipoproteins”, “Metabolism”, “Signaling by Rho GTPases” show in both paternal as well as maternal TRD regions. This provides support for the hypothesis that fundamental cell cycle and metabolic processes are affected by TRD and that selection of major phenotypic differences (e.g. body weight and fat composition) shaped the allelic composition of the genome of the founder inbred lines by different genetic requirements. These genetic adaptations are necessary for the optimization of the genome to ensure fitness and reproduction during the generation of inbred lines and might be what causes the observed TRD when founder genomes are combined together.

Our study sheds new light on the TRD in a cross between different inbred mouse strains, the distinct functioning of genomes in producing viable offspring, and provides a way to identify candidate genes which could contribute to complex traits different between the founder strains (in our case obesity and/or retinal functionality). The genes in the TRD regions provide new targets for investigating genetic adaptation and modifying determinants of complex traits.

## Materials and Methods

### Mouse population

348 male mice of an advanced intercross line (AIL) in generation 28 as well as their 62 parents and 66 grandparents from generations 27 and 26 were genotyped. The AIL population originates from the mapping population of a cross between a male mouse of the obese line BFMI860-12 (BFMI) and a female of the lean line C57BL/6NCrl (B6N), that had been initially used to map the juvenile obesity locus *jObes1* (Neuschl et al. 2010). Beginning in generation F_1_, individuals were randomly mated to mice from the same generation using the program RandoMate (Schmitt et al. 2009).

### Husbandry conditions

All experimental treatments of animals were approved by the German Animal Welfare Authorities (approval no. G0016/11). All mice were maintained under conventional conditions and a 12:12 h light:dark cycle (lights on at 6:00 am) at a temperature of 22 ± 2 °C. Animals had *ad libitum* access to food and water. To perform fine mapping of the obesity QTL (Arends et al. 2016), generation 28 was fed with a rodent high-fat diet (HFD) containing 19.5 MJ/kg of metabolizable energy, 45% from fat, 24% from protein and 31% from carbohydrates (E15103-34, ssniff EF R/M, Ssniff Spezialdiäten GmbH, Soest / Germany). All other generations used in this study were fed a standard breeding diet (V1534-000, ssniff EF R/M; Ssniff Spezialdiäten GmbH, Soest, Germany).

### Genotypes

Genotypes were generated at GeneSeek (Lincoln, NE, USA) using the Mega Mouse Universal Genotyping Array (MegaMUGA). These arrays are SNP genotyping arrays based on the Illumina Infinium platform designed by investigators at the University of North Carolina at Chapel Hill, manufactured by Illumina (San Diego, CA), and distributed by Neogen Inc (Lansing, MI, USA) (Morgan and Welsh 2015). This array contains probes targeting 77,800 known SNPs. SNP probes were remapped to the reference genome (GRCm38_95) using BLASTN with default settings (Camacho et al. 2009). In order to increase the certainty of genotype calls, genotypes with a GenCall score greater than 0.7 were considered confidently called, although the manufacturer’s recommendation is a GenCall score > 0.15. SNPs that mapped to multiple positions in the genome, non-informative SNPs, and SNPs with genotype call rates below 90% were removed from further analysis. In total, 14,415 highly confident SNPs passed all quality checks and were informative between BFMI and B6N. Marker density, as well as minor allele frequencies (MAF) within and outside of TRD regions were visualized and can be found in Supplemental_Table_S5.

Furthermore, checking parent-child relations in our trio data identified 3 individuals in generation 28 where one of the parents was wrongly assigned, these 3 individuals were removed from further analysis. Similarly, one individual in generation 27 was found to have a wrong parent assignment, leading to the removal of this individual and its 4 offspring in generation 28. Phasing of the heterozygous genotypes of the AIL animals of generation 28 towards the parental population (generation 27), and of generation 27 to generation 26 was done using Beagle v4.1 (Browning and Browning 2007) with standard settings. Raw and phased genotypes of all individuals that passed quality control (N_(28)_ = 341, N_(27)_ = 61, and N_(26)_ = 66), the genetic map, and pedigree data are available in Supplemental_Table_S2.

### Allele transmission from heterozygous parents

Deviations from expected Mendelian inheritance ratios are named transmission ratio distortion (TRD). Such deviations have been commonly observed in experimental crosses as well as in natural populations. We used an extension of the transmission asymmetry test and parental asymmetry test to detect parent-of-origin dependent effects on the frequency of the transmission of a specific SNP allele from parent to offspring using trios in our AIL design (Weinberg et al. 1998). For example: To determine if one of the alternative paternal alleles (e.g. A versus B allele) at a SNP locus is inherited more often than expected by Mendel (50%), pups were analyzed in generation 28 of fathers (generation 27) that were heterozygous for this SNP. We only tested markers at which at least 10 heterozygous fathers (or mothers) were available. We counted the number of offspring where a specific paternal allele was transmitted. When both parents were heterozygous, the allele transmitted cannot be determined and this transmission was not counted in the test statistic. Furthermore, markers were tested for Hardy Weinberg equilibrium (HWE) using the code developed by Wigginton *et al.* (Wigginton et al. 2005). Markers not in HWE were excluded, since the χ^2^ test for TRD is only valid when a marker is in HWE. A χ^2^ test was used to test if this distribution of paternally inherited alleles significantly deviated from the expected Mendelian inheritance ratios (**Pat**), and similarly for maternally inherited alleles (**Mat**).

- ***Pat***: Analysis of allele transmission ratio distortion from heterozygous fathers to offspring

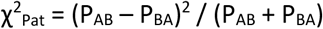
- ***Mat***: Analysis of allele transmission ratio distortion from heterozygous mothers to offspring

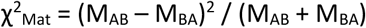

*χ*^2^ scores were transformed into p-values using the appropriate conversions and then transformed into LOD scores using –log_10_(p-value). 5% and 1% significance thresholds were determined by Bonferroni correction (5% >= 6.75, 1% >= 7.45). Significant regions were defined as the region from the first to the last flanking marker above the 1% significance threshold (LOD scores >= 7.45).

### Genetic variants in TRD regions

Parental genomes (BFMI860-12 and B6N) were paired-end sequenced using the “Illumina HiSeq” platform (Illumina Inc., San Diego, California, U.S.). Obtained DNA reads were trimmed using trimmomatic (Bolger et al. 2014) after which trimmed reads were aligned to the mouse genome (MM10, GRCm38.p6) using the Burrows–Wheeler Aligner (BWA) software (Li and Durbin 2009). The subsequent SAM files were converted to BAM files, sorted, and indexed using Samtools (Li et al. 2009; Morgan et al. 2017). (Optical) Duplicate reads were removed using Picard tools v2.19.0 (Broad Institute 2016), after which indel realignment and base recalibration was done using the GATK v4.1.0.0 (McKenna et al. 2010), according to GATK best practices (Broad Institute 2018). Sequence variants were called using BCFtools (Morgan et al. 2017) Variants passing quality control were further annotated using the Ensembl Variant Effect Predictor (VEP) (McLaren et al. 2016). DNA sequencing data allowed to identify non-synonymous SNPs (nsSNPs) in genes located in TRD regions between the founding strains.

A permutation strategy was used to detect over- and/or underrepresentation of nsSNPs in the regions showing TRD. We performed 50,000 permutations, each time drawing 1,424 protein-coding genes at random, not allowing duplicate genes or selection of predicted genes (GM/RIKEN). For every permutation, the number of nsSNPs and the number of genes with nsSNPs was recorded. After 50,000 permutations, a distribution of the total number of nsSNPs (and genes) in the random data was obtained, which was compared with the observed data.

### Pathway overrepresentation analyses

We extracted all protein-coding genes inside the significant regions using biomaRt (Kasprzyk 2011) for each of the different types of allele transmission ratio distortion: Preferred paternal transmission of the BFMI allele (Pat_BFMI), preferred paternal transmission of the B6N allele (Pat_B6N), preferred maternal transmission of the BFMI allele (Mat_BFMI), and preferred maternal transmission of the B6N allele (Mat_B6N). To identify potential functional clustering of genes within one of these groups, pathway overrepresentation analyses was performed using innateDB (Breuer et al. 2013) with KEGG (Ogata et al. 2000) and Reactome (Joshi-Tope et al. 2005) as the pathway providers. Overrepresentation was tested using a hypergeometric test. P-values reported for pathway overrepresentation were Benjamini-Hochberg corrected (P_(BH)_) (Benjamini and Hochberg 1995); P_(BH)_ <0.05 were considered significant. Pathway analysis was additionally performed with genes showing nsSNPs using the same grouping as before, with “_SNP” added to the group identifier (Pat_BFMI_SNP, Mat_BFMI_SNP, Pat_B6N_SNP, and Mat_B6N_SNP).

### Genetic incompatibilities

Testing for pairwise genetic incompatibilities in an exhaustive manner is not advisable because of the large number of statistical tests required for 20k SNP markers leading to a severe multiple testing correction. Our hypothesis is that genetic incompatibilities cause allele transmission ratio distortions. For testing incompatibilities, 3×3 contingency tables were created using the top SNP marker in the TRD region 1 (M1) versus the top SNP marker in TRD region 2 (M1), and the number of co-occurrences between different alleles was counted. If no top marker was present in a region e.g. Pat_R5, due to all markers showing a similar distortion, the proximal flanking marker was used as top marker. Our method for scoring genetic incompatibility is very similar to the methods used by Ackermann and Beyer and Corbett-Detig et al (Ackermann and Beyer 2012; Corbett-Detig et al. 2013). A 3×3 table of expected co-occurrences based on the observed allele frequencies at markers M1 and M2 was generated assuming independent segregation of each marker. Resulting *χ*^2^ scores were transformed into p-values, which are then transformed to LOD scores as described before. For each pair of markers that showed a genome-wide significant interaction (P_(BF)_ < 0.05), founder alleles of the group which shows the most reduction (in percentage) between observed and expected co-occurrences was used for the visualization seen in Figure 2.

Pairwise interactions tests were only performed between detected TRD regions, while correction for multiple testing was done using genome-wide thresholds. This first involved estimating the number of effective tests by using the simpleM method (Gao 2011). The simpleM method was designed to estimate the number of independent tests in a GWAS by considering linkage between markers. The simpleM procedure estimated 1,008 independent test (at a fixLength of 1200) which is much lower than the number of genetic markers. This reduction in total tests can be explained by strong linkage between markers in our AIL population. LOD thresholds were adjusted for multiple testing using Bonferroni correction (P_(BF)_) and the number of independent tests estimated with the simpleM method (n=1,008). Since we tested pairwise but without repeating the test for a pair we have already tested, the number of tests had to be multiplied with itself and reduced by half leading to LOD scores calculated as: −log10(threshold / (1,008 * 1,008 * 0.5). Dependent on the significance threshold, this leads to the following genome-wide adjusted LOD thresholds: significant if LOD > 7.0 (P_(BF)_ < 0.05) and highly significant if LOD > 7.7 (P_(BF)_ < 0.01).

We then continued our investigation of known protein-protein interactions between genes with non-synonymous SNPs (nsSNPs) within these regions of incompatibility using the Search Tool for the Retrieval of Interacting Genes/Proteins (STRING) database version 11 (Szklarczyk et al. 2019). In total 9,602,772 known physical protein-protein interactions for *Mus musculus* (SpeciesID 10090) are listed in this database (10090.protein.physical.links.v11.0). We first only considered the protein-protein interactions between the 128 genes with one or more nsSNP(s). Afterwards, we overlaid the gene location data with the TRD regions for which we found genome-wide significant evidence of genetic incompatibilities. This was done to see if identified genetic compatibility could be explained by known physical protein-protein interactions in which both participating genes show one or more nsSNP(s).

## List of abbreviations

AIL: Advanced intercross line
BFMI: Berlin Fat Mouse inbred line
B6N: C57BL/6N inbred line
Mat: Maternal transmission ratio distortion
MAF: Minor allele frequency
nsSNP: Non-Synonymous Single Nucleotide Polymorphism
P_(BF)_: Bonferroni corrected p-value
P_(BH)_: Benjamini–Hochberg corrected p-value
Pat: Paternal transmission ratio distortion
QTL: Quantitative trait locus
SNP: Single nucleotide polymorphism
TRD: Transmission ratio distortion

## Declarations

### Ethics approval

All experimental treatments of animals were approved by the German Animal Welfare Authorities (approval no. G0016/11).

### Availability of data and materials

The datasets supporting the conclusions of this article are included within the article and its supplementary files. DNA sequencing data was deposited at the NCBI Sequence Read Archive (SRA) under BioProject ID: PRJNA717237

### Authors’ contributions

**D.A.** contributed to the design of the experiment, performed the data analysis and drafted the manuscript. **S. K.** was involved in the design of the experiment, animal handling, sample collection, and contributed to the manuscript. **S.H.** was responsible for animal handling, sample collection and DNA extraction. **P.K.** contributed to critical data analysis discussions, revision of the manuscript and Table 1, and Figure 2. **D.H.** contributed to writing and critical revision of the manuscript, and discussion about the biological implications of the observed TRD. **G.A.B.** designed the experiment, planned, organized and supervised the project; contributed to write and critically revise the manuscript.

### Authors’ information

Dr. Danny Arends finished his PhD at the Groningen Bioinformatics Centre at the University of Groningen in November 2014 and is currently working as a PostDoc at the Humboldt University in Berlin. He develops new computational methodologies and open source software for use in (systems) biology research, such as the addition of Multiple QTL Mapping (MQM) into R/qtl. His focus is development of large-scale computational methods in QTL analysis, and the integration of ‘big data’ biological data sets in a comprehensive manner.

Dr. Sebastian Heise finished his PhD in the group Breeding Biology and molecular Genetics at Humboldt-University in Berlin in November 2014. He is working with lean and obese mouse models on projects focusing on insulin sensitivity and type-2 diabetes. Exploring the genetics contribution beyond the complex traits is the key interest of his projects.

Dr. Stefan Kärst finished his PhD in the group Breeding Biology and molecular Genetics at Humboldt-University in Berlin in October 2011 and is currently working at a Biotech company in Berlin. During his PhD and PostDoc at the Humboldt University in Berlin, Stefan developed new methods for phenotyping, mapped muscle trait QTLs and analyzed NGS data for gene expression and sequence variants.

Dr. Paula Korkuc obtained her Bioinformatics Ph.D. degree from the University of Potsdam in cooperation with the Max-Planck institute for Molecular Plant Physiology in 2016. She is currently working as a PostDoc at the Humboldt University in Berlin. Her current research focusses on the identification of functional variants associated with milk production traits in the endangered German DSN cattle breed using DNA sequencing, SNP-chip data, and novel computational methods implemented in R.

Dr. Deike Hesse studied biology at the universities of Düsseldorf, Nantes and Marburg. She obtained her doctor degree from the University of Potsdam in collaboration with the German Institute for Human Nutrition in 2010. Since 2017 she works as a researcher and lecturer at the Humboldt University in Berlin. Her research focusses on energy metabolism, body weight development, and diabetes. Using mouse models with different genetic backgrounds she aims to elucidate the contribution and interaction of different tissues and metabolic pathways to metabolic phenotypes.

Prof. Dr. Gudrun A. Brockmann studied Biophysics in Berlin, finished her PhD in Embryology at the Academy of Agricultural Sciences, and obtained her Habilitation in the field of Genetics at Rostock University, Germany. She had worked with mice as models for animal breeding at the Leibnitz-Institute for Farm Animals in Dummerstorf, before she was appointed as a Full Professor for Animal Breeding and Molecular Genetics at Humboldt-University Berlin in 2003. Her interest is the understanding of genetics underlying complex traits with a focus on energy partitioning. She is author of more than 130 papers, acts on scientific advisory boards, and as reviewer for all kind of research applications.

## Supplemental files

Supplemental_Table_S1 - ***Transmission ratio distortion, allele transmissions, and p-values***

Supplemental_Table_S2 – ***Genotypes, map, and pedigree of the AIL individuals***

Supplemental_Table_S3 - ***Genes in TRD regions (All, Non-synonymous SNPs)***

Supplemental_Table_S4 - **Complete *pathway over-representation analysis***

Supplemental_File_S5 - **MAF comparison and marker density**

## References

Ackermann M, Beyer A. 2012. Systematic Detection of Epistatic Interactions Based on Allele Pair Frequencies ed. J. Flint. PLoS Genet 8: e1002463. https://dx.plos.org/10.1371/journal.pgen.1002463.

Arends D, Heise S, Kärst S, Trost J, Brockmann GA. 2016. Fine mapping a major obesity locus (jObes1) using a Berlin Fat Mouse × B6N advanced intercross population. Int J Obes 40: 1784–1788. http://www.nature.com/articles/ijo2016150.

Benjamini Y, Hochberg Y. 1995. Controlling the False Discovery Rate: A Practical and Powerful Approach to Multiple Testing. J R Stat Soc Ser B 57: 289–300. http://www.jstor.org/stable/pdfplus/2346101.pdf?acceptTC=true.

Blake JA, Baldarelli R, Kadin JA, Richardson JE, Smith CL, Bult CJ, Anagnostopoulos A V, Beal JS, Bello SM, Blodgett O, et al. 2021. Mouse Genome Database (MGD): Knowledgebase for mouse– human comparative biology. Nucleic Acids Res 49: D981–D987. https://academic.oup.com/nar/article/49/D1/D981/5999894.

Bolger AM, Lohse M, Usadel B. 2014. Trimmomatic: a flexible trimmer for Illumina sequence data. Bioinformatics 30: 2114–2120. http://bioinformatics.oxfordjournals.org/content/30/15/2114%5Cnhttp://bioinformatics.oxfordjournals.org/content/30/15/2114.abstract.html?etoc%5Cnhttp://bioinformatics.oxfordjournals.org/content/30/15/2114.full.pdf%5Cnhttp://www.ncbi.nlm.nih.gov/pubmed/24695404 (Accessed July 9, 2014).

Brand CL, Larracuente AM, Presgraves DC. 2015. Origin, evolution, and population genetics of the selfish Segregation Distorter gene duplication in European and African populations of Drosophila melanogaster. Evolution (N Y).

Breuer K, Foroushani AK, Laird MR, Chen C, Sribnaia A, Lo R, Winsor GL, Hancock REW, Brinkman FSL, Lynn DJ. 2013. InnateDB: Systems biology of innate immunity and beyond - Recent updates and continuing curation. Nucleic Acids Res 41.

Broad Institute. 2018. GATK Best Practices. https://software.broadinstitute.org/gatk/best-practices/ (Accessed February 1, 2018).

Broad Institute. 2016. Picard tools. https://broadinstitute.github.io/picard/.

Brockmann C., Brockmann T., Dege S., Skosyrski S., Strauß O, Joussen A. M. 2017. SaF08-04 - Hereditäre Retinale Degeneration der Berliner Fettmaus. In Abstractband DOG, p. 201 https://www.dog.org/wp-content/uploads/2017/08/Abstractband_DOG_2017.pdf.

Browning SR, Browning BL. 2007. Rapid and accurate haplotype phasing and missing-data inference for whole-genome association studies by use of localized haplotype clustering. Am J Hum Genet 81: 1084–1097.

Brunet S, Verlhac MH. 2011. Positioning to get out of meiosis: The asymmetry of division. Hum Reprod Update.

Buniello A, MacArthur JAL, Cerezo M, Harris LW, Hayhurst J, Malangone C, McMahon A, Morales J, Mountjoy E, Sollis E, et al. 2019. The NHGRI-EBI GWAS Catalog of published genome-wide association studies, targeted arrays and summary statistics 2019. Nucleic Acids Res 47: D1005–D1012. https://academic.oup.com/nar/article/47/D1/D1005/5184712.

Camacho C, Coulouris G, Avagyan V, Ma N, Papadopoulos J, Bealer K, Madden TL. 2009. BLAST+: architecture and applications. BMC Bioinformatics 10: 421.

Chmátal L, Gabriel SI, Mitsainas GP, Martínez-Vargas J, Ventura J, Searle JB, Schultz RM, Lampson MA. 2014. Centromere strength provides the cell biological basis for meiotic drive and karyotype evolution in mice. Curr Biol.

Clark GW, Dar V-N, Bezginov A, Yang JM, Charlebois RL, Tillier ERM. 2011. Using Coevolution to Predict Protein–Protein Interactions. pp. 237–256 http://link.springer.com/10.1007/978-1-61779-276-2_11.

Comuzzie AG, Cole SA, Laston SL, Voruganti VS, Haack K, Gibbs RA, Butte NF. 2012. Novel Genetic Loci Identified for the Pathophysiology of Childhood Obesity in the Hispanic Population. PLoS One.

Corbett-Detig RB, Zhou J, Clark AG, Hartl DL, Ayroles JF. 2013. Genetic incompatibilities are widespread within species. Nature 504: 135–7. http://www.ncbi.nlm.nih.gov/pubmed/24196712.

DeBosch BJ, Kluth O, Fujiwara H, Schürmann A, Moley K. 2014. Early-onset metabolic syndrome in mice lacking the intestinal uric acid transporter SLC2A9. Nat Commun 5: 4642. http://www.nature.com/articles/ncomms5642.

Dinour D, Gray NK, Campbell S, Shu X, Sawyer L, Richardson W, Rechavi G, Amariglio N, Ganon L, Sela BA, et al. 2010. Homozygous SLC2A9 mutations cause severe renal hypouricemia. J Am Soc Nephrol.

Duriez B, Duquesnoy P, Escudier E, Bridoux AM, Escalier D, Rayet I, Marcos E, Vojtek AM, Bercher JF, Amselem S. 2007. A common variant in combination with a nonsense mutation in a member of the thioredoxin family causes primary ciliary dyskinesia. Proc Natl Acad Sci U S A.

Fel-Clair F, Catalan J, Lenormand T, Britton-Davidian J. 1998. Centromeric Incompatibilities in the Hybrid Zone Between House Mouse Subspecies from Denmark: Evidence from Patterns of Nor Activity. Evolution (N Y) 52: 592–603.

Fischer E, Charbonneau H, Tonks N. 1991. Protein tyrosine phosphatases: a diverse family of intracellular and transmembrane enzymes. Science (80−) 253: 401–406. https://www.sciencemag.org/lookup/doi/10.1126/science.1650499.

Fitzpatrick JL, Evans JP. 2009. Reduced heterozygosity impairs sperm quality in endangered mammals. Biol Lett 5: 320–323. https://royalsocietypublishing.org/doi/10.1098/rsbl.2008.0734.

Flurkey K, Currer JM, Leiter EH, Witham B, Jackson Laboratory (Bar Harbor M. 2009. The Jackson Laboratory Handbook on Genetically Standardized Mice. 6th ed. eds. K. Flurkey, J.M. Currer, E.H. Leiter, and B. Witham. The Jackson Laboratory, Bar Harbor, ME 04609 USA https://www.urmc.rochester.edu/MediaLibraries/URMCMedia/animal-resource/forms/documents/JAX-Handbook-Genetically-Standardized-Mice.pdf.

Gao X. 2011. Multiple testing corrections for imputed SNPs. Genet Epidemiol 35: 154–158.

GeneCards Human Gene Database. 2021. STK32B Gene - GeneCards. https://www.genecards.org/cgi-bin/carddisp.pl?gene=STK32B.

Heid IM, Jackson AU, Randall JC, Winkler TW, Qi L, Steinthorsdottir V, Thorleifsson G, Zillikens MC, Speliotes EK, Mägi R, et al. 2010. Meta-analysis identifies 13 new loci associated with waist-hip ratio and reveals sexual dimorphism in the genetic basis of fat distribution. Nat Genet 42: 949–960. https://www.nature.com/articles/ng.686.pdf.

Heinonen S, Buzkova J, Muniandy M, Kaksonen R, Ollikainen M, Ismail K, Hakkarainen A, Lundbom J, Lundbom N, Vuolteenaho K, et al. 2015. Impaired mitochondrial biogenesis in adipose tissue in acquired obesity. Diabetes.

Hirsch CN, Flint-Garcia SA, Beissinger TM, Eichten SR, Deshpande S, Barry K, McMullen MD, Holland JB, Buckler ES, Springer N, et al. 2014. Insights into the Effects of Long-Term Artificial Selection on Seed Size in Maize. Genetics 198: 409–421. https://academic.oup.com/genetics/article/198/1/409-421/6073421.

Huang LO, Labbe A, Infante-Rivard C. 2013a. Transmission ratio distortion: review of concept and implications for genetic association studies. Hum Genet 132: 245–63. http://www.ncbi.nlm.nih.gov/pubmed/23242375.

Huang X, Baumann M, Nikitina L, Wenger F, Surbek D, Körner M, Albrecht C. 2013b. RNA degradation differentially affects quantitative mRNA measurements of endogenous reference genes in human placenta. Placenta 34: 544–7. http://www.ncbi.nlm.nih.gov/pubmed/23623484. (Accessed October 11, 2013)

Joshi-Tope G, Gillespie M, Vastrik I, D’Eustachio P, Schmidt E, de Bono B, Jassal B, Gopinath GR, Wu GR, Matthews L, et al. 2005. Reactome: a knowledgebase of biological pathways. Nucleic Acids Res 33: D428–32. http://www.ncbi.nlm.nih.gov/pubmed/15608231.

Kasprzyk A. 2011. BioMart: driving a paradigm change in biological data management. Database (Oxford) 2011: bar049. http://www.pubmedcentral.nih.gov/articlerender.fcgi?artid=3215098&tool=pmcentrez&rendertype=abstract. (Accessed July 12, 2014)

Klink BU, Gatsogiannis C, Hofnagel O, Wittinghofer A, Raunser S. 2020. Structure of the human BBSome core complex. Elife 9. https://elifesciences.org/articles/53910.

Kopelman P. 2007. Health risks associated with overweight and obesity. Obes Rev 8: 13–17. http://doi.wiley.com/10.1111/j.1467-789X.2007.00311.x.

Kursel LE, Malik HS. 2018. The cellular mechanisms and consequences of centromere drive. Curr Opin Cell Biol.

Lanneluc I, Desmarais E, Boursot P, Dod B, Bonhomme F. 2004. Characterization of a centromeric marker on mouse Chromosome 11 and its introgression in a domesticus/musculus hybrid zone. Mamm Genome.

Larson EL, Vanderpool D, Sarver BAJ, Callahan C, Keeble S, Provencio LL, Kessler MD, Stewart V, Nordquist E, Dean MD, et al. 2018. The evolution of polymorphic hybrid incompatibilities in house mice. Genetics.

Lenormand T, Fel-Clair F, Manolakou K, Alibert P, Britton-Davidian J. 1997. Chromosomal transmission bias in laboratory hybrids between wild strains of the two European subspecies of house mice. Genetics.

Li G, Jin J, Zhou Y, Bai X, Mao D, Tan C, Wang G, Ouyang Y. 2019. Genome-wide dissection of segregation distortion using multiple inter-subspecific crosses in rice. Sci China Life Sci 62: 507–516. http://link.springer.com/10.1007/s11427-018-9452-8.

Li H, Durbin R. 2009. Fast and accurate short read alignment with Burrows-Wheeler transform. Bioinformatics 25: 1754–60. https://www.ncbi.nlm.nih.gov/pmc/articles/PMC2705234/. (Accessed July 9, 2014)

Li H, Handsaker B, Wysoker A, Fennell T, Ruan J, Homer N, Marth G, Abecasis GR, Durbin R. 2009. The Sequence Alignment/Map format and SAMtools. Bioinformatics 25: 2078–2079. https://academic.oup.com/bioinformatics/article-lookup/doi/10.1093/bioinformatics/btp352.

Locke AE, Kahali B, Berndt SI, Justice AE, Pers TH, Day FR, Powell C, Vedantam S, Buchkovich ML, Yang J, et al. 2015. Genetic studies of body mass index yield new insights for obesity biology. Nature 518: 197–206. http://dx.doi.org/10.1038/nature14177.

Lu M, Varley AW, Ohta S, Hardwick J, Munford RS. 2008. Host Inactivation of Bacterial Lipopolysaccharide Prevents Prolonged Tolerance Following Gram-Negative Bacterial Infection. Cell Host Microbe 4: 293–302. https://linkinghub.elsevier.com/retrieve/pii/S1931312808002242.

Lyon MF. 2003. Transmission ratio distortion in mice. Annu Rev Genet 37: 393–408. http://www.ncbi.nlm.nih.gov/pubmed/14616067.

Martin-DeLeon PA, Zhang H, Morales CR, Zhao Y, Rulon M, Barnoski BL, Chen H, Galileo DS. 2005. Spam I-associated transmission ratio distortion in mice: Elucidating the mechanism. Reprod Biol Endocrinol.

Matsuo H, Chiba T, Nagamori S, Nakayama A, Domoto H, Phetdee K, Wiriyasermkul P, Kikuchi Y, Oda T, Nishiyama J, et al. 2008. Mutations in Glucose Transporter 9 Gene SLC2A9 Cause Renal Hypouricemia. Am J Hum Genet.

McKenna A, Hanna M, Banks E, Sivachenko AY, Cibulskis K, Kernytsky AM, Garimella K, Altshuler D, Gabriel S, Daly MJ, et al. 2010. The Genome Analysis Toolkit: A MapReduce framework for analyzing next-generation DNA sequencing data. Genome Res 20: 1297–1303. http://genome.cshlp.org/cgi/doi/10.1101/gr.107524.110. (Accessed July 9, 2014)

McLaren W, Gil L, Hunt SE, Riat HS, Ritchie GRS, Thormann A, Flicek P, Cunningham F. 2016. The Ensembl Variant Effect Predictor. Genome Biol 17: 1–14.

Morgan AP, Welsh CE. 2015. Informatics resources for the Collaborative Cross and related mouse populations. Mamm Genome.

Morgan M, Pagès H, Obenchain V, N H. 2017. Rsamtools: Binary alignment (BAM), FASTA, variant call (BCF), and tabix file import. R Packag version 1300. http://bioconductor.org/packages/release/bioc/html/Rsamtools.html. (Accessed February 15, 2018)

Mouse Genome Database (MGD). 2021. Myo5c - MGI Mouse Gene Detail. http://www.informatics.jax.org/marker/MGI:2442485 (Accessed September 3, 2021).

Neuschl C, Hantschel C, Wagener A, Schmitt AO, Illig T, Brockmann GA. 2010. A unique genetic defect on chromosome 3 is responsible for juvenile obesity in the Berlin Fat Mouse. Int J Obes 34: 1706–1714. https://www.nature.com/articles/ijo201097.

Ogata H, Goto S, Sato K, Fujibuchi W, Bono H, Kanehisa M, Goto S. 2000. KEGG: kyoto encyclopedia of genes and genomes. Nucleic Acids Res 28: 29–34. https://www.ncbi.nlm.nih.gov/pmc/articles/PMC102409/.

Ojogun NI. 2008. CONSTITUTIVE OVEREXPRESSION OF ACYLOXYACYL HYDROLASE IN MUS MUSCULUS. The University of Texas Southwestern Medical Center at Dallas https://utswmed-ir.tdl.org/bitstream/handle/2152.5/383/OjogunNoredia.pdf.

Papes F, Kemper EL, Cord-Neto G, Langone F, Arruda P. 1999. Lysine degradation through the saccharopine pathway in mammals: involvement of both bifunctional and monofunctional lysine-degrading enzymes in mouse. Biochem J 344 Pt 2: 555–63. https://portlandpress.com/biochemj/article/344/2/555/36179/Lysine-degradation-through-the-saccharopine.

Paterson AD, Waggott D, Schillert A, Infante-Rivard C, Bull SB, Yoo YJ, Pinnaduwage D. 2009. Transmission-ratio distortion in the Framingham Heart Study. BMC Proc 3: S51. https://bmcproc.biomedcentral.com/articles/10.1186/1753-6561-3-S7-S51.

Rodríguez-Carrizalez AD, Castellanos-González JA, Martínez-Romero EC, Miller-Arrevillaga G, Villa-Hernández D, Hernández-Godínez PP, Ortiz GG, Pacheco-Moisés FP, Cardona-Muñoz EG, Miranda-Díaz AG. 2014. Oxidants, antioxidants and mitochondrial function in non-proliferative diabetic retinopathy. J Diabetes.

Rodriguez OC, Cheney RE. 2002. Human myosin-Vc is a novel class V myosin expressed in epithelial cells. J Cell Sci.

Sacksteder KA, Biery BJ, Morrell JC, Goodman BK, Geisbrecht B V., Cox RP, Gould SJ, Geraghty MT. 2000. Identification of the α-aminoadipic semialdehyde synthase which is defective in familial hyperlysinemia. Am J Hum Genet.

Safronova LD, Chubykin VL. 2013. Meiotic drive in mice carrying t-complex in their genome. Russ J Genet.

Schmitt AO, Bortfeldt RH, Neuschl C, Brockmann GA. 2009. RandoMate: A program for the generation of random mating schemes for small laboratory animals. Mamm Genome 20: 321–325.

Shungin D, Winkler TW, Croteau-Chonka DC, Ferreira T, Locke AE, Mägi R, Strawbridge RJ, Pers TH, Fischer K, Justice AE, et al. 2015. New genetic loci link adipose and insulin biology to body fat distribution. Nature 518: 187–96. http://dx.doi.org/10.1038/nature14132.

Singhai M, Goyal R, Faizy A. 2011. Glutathione peroxidase activity in obese and nonobese diabetic patients and role of hyperglycemia in oxidative stress. J Midlife Health.

Speliotes EK, Willer CJ, Berndt SI, Monda KL, Thorleifsson G, Jackson AU, Allen HL, Lindgren CM, Luan J, Mägi R, et al. 2010. Association analyses of 249,796 individuals reveal 18 new loci associated with body mass index. Nat Genet 42: 937–948. https://www.nature.com/articles/ng.686.pdf.

Szklarczyk D, Gable AL, Lyon D, Junge A, Wyder S, Huerta-Cepas J, Simonovic M, Doncheva NT, Morris JH, Bork P, et al. 2019. STRING v11: Protein-protein association networks with increased coverage, supporting functional discovery in genome-wide experimental datasets. Nucleic Acids Res.

Teeter KC, Payseur BA, Harris LW, Bakewell MA, Thibodeau LM, O’Brien JE, Krenz JG, Sans-Fuentes MA, Nachman MW, Tucker PK. 2008. Genome-wide patterns of gene flow across a house mouse hybrid zone. Genome Res 18: 67–76.

Temtamy SA, Aglan MS, Valencia M, Cocchi G, Pacheco M, Ashour AM, Amr KS, Helmy SMH, El-Gammal MA, Wright M, et al. 2008. Long interspersed nuclear element-1 (LINE1)-mediated deletion of EVC, EVC2, C4orf6, and STK32B in ellis-van Creveld syndrome with borderline intelligence. Hum Mutat.

Teppa E, Zea DJ, Marino-Buslje C. 2017. Protein–protein interactions leave evolutionary footprints: High molecular coevolution at the core of interfaces. Protein Sci.

Tremmel M, Gerdtham U-G, Nilsson P, Saha S. 2017. Economic Burden of Obesity: A Systematic Literature Review. Int J Environ Res Public Health 14: 435. http://www.mdpi.com/1660-4601/14/4/435.

Wagener A, Schmitt AO, Aksu S, Schlote W, Neuschl C, Brockmann GA. 2006. Genetic, sex, and diet effects on body weight and obesity in the Berlin Fat Mouse Inbred lines. Physiol Genomics 27: 264–270.

Weinberg CR, Wilcox AJ, Lie RT. 1998. A log-linear approach to case-parent-triad data: assessing effects of disease genes that act either directly or through maternal effects and that may be subject to parental imprinting. Am J Hum Genet 62: 969–978.

Whitlock MC. 2000. Fixation Of New Alleles And The Extinction Of Small Populations: Drift Load, Beneficial Alleles, And Sexual Selection. Evolution (N Y) 54: 1855–1861. http://doi.wiley.com/10.1111/j.0014-3820.2000.tb01232.x.

Wigginton JE, Cutler DJ, Abecasis GR. 2005. A Note on Exact Tests of Hardy-Weinberg Equilibrium. Am J Hum Genet 76: 887–893. https://linkinghub.elsevier.com/retrieve/pii/S0002929707607356.

Willer CJ, Speliotes EK, Loos RJF, Li S, Lindgren CM, Heid IM, Berndt SI, Elliott AL, Jackson AU, Lamina C, et al. 2009. Six new loci associated with body mass index highlight a neuronal influence on body weight regulation. Nat Genet 41: 25–34. http://www.nature.com/doifinder/10.1038/ng.287.

Xie Y, Tang J, Xie X, Li X, Huang J, Fei Y, Han J, Chen S, Tang H, Zhao X, et al. 2019. An asymmetric allelic interaction drives allele transmission bias in interspecific rice hybrids. Nat Commun.

Zajitschek SRK, Lindholm AK, Evans JP, Brooks RC. 2009. Experimental evidence that high levels of inbreeding depress sperm competitiveness. J Evol Biol 22: 1338–45. http://doi.wiley.com/10.1111/j.1420-9101.2009.01738.x.

Zuk O, Hechter E, Sunyaev SR, Lander ES. 2012. The mystery of missing heritability: Genetic interactions create phantom heritability. Proc Natl Acad Sci 109: 1193–1198. http://www.pnas.org/cgi/doi/10.1073/pnas.1119675109.

